# Effects of High-Intensity Interval Training on Physical and Cognitive Function in Middle-Aged Male Mice

**DOI:** 10.1101/2025.02.13.638126

**Authors:** Justin C. Stephenson, Tuan D. Tran, Ted G. Graber

**Author notes:** Corresponding/Senior Author: Ted G. Graber, PhD., ECU College of Allied Health Sciences Department of Physical Therapy, Health Sciences Building, Rm 2410 | Mail Stop 668 Greenville, NC 27834.

## Abstract

Declining functional capacity, both physical and cognitive, is a consequence of aging. However, exercise is a promising intervention to mitigate normal age-related decline. While numerous studies have elucidated the benefits of exercise per se, less well-studied is the effect of high intensity interval training (HIIT) on a middle-aged population. Our primary purpose was to assess the effect of three months of HIIT on physical and cognitive performance in middle-aged (17-month-old) male C57BL/6J mice, compared to sedentary controls. We hypothesized that exercised mice would be resistant to any decline in cognitive and physical ability, both measured pre- and post-intervention. To measure physical function, we used the well-validated CFAB (comprehensive functional assessment battery) scoring system comprised of determinants including voluntary wheel running, inverted cling, grip test, treadmill max speed, and rotarod. We measured cognition with open field, novel object recognition, y-maze, and puzzle box. Further measures of sarcopenia/frailty included body composition (MRI) and *in vivo* contractile physiology (plantar flexor torque). Training resulted in significant aerobic capacity improvements for the HIIT group, increasing treadmill time by 28%, while the SED group demonstrated a 41.4% decline in treadmill time. However, no significant differences in cognitive function were determined. Contrary to our previous research in other age groups, the current study found a negligible effect of HIIT on body composition. We note that at 17 months old, mice did not experience any evidence of cognitive deterioration in either group over the three-month period, thus explaining the lack of exercise effect. We found that HIIT had less influence on either physical or cognitive function than we expected, which may be because function in this age group is stable. Future work will investigate older adult cognitive response to HITT at ages where there is well-documented cognitive decline.

## INTRODUCTION

In the United States alone, the percentage of adults ≥ 65 years of age increased 38.6% between 2010 and 2020.^1^ Compared to 2015, the percentage of adults ≥ 60 years of age, globally, is projected to reach ≥ 16.5% by 2030, marking an increase of more than 4% within only 15 years.^2^ Such developments and age distribution shifts toward an older population will increase the prevalence of age-related functional (physical and cognitive) declines, as well as increased need for medical, residential, and home care.

As we grow older, mild changes in cognition are expected and considered a normal feature of the aging process,^3^ also known as *normal cognitive decline*. Evidence suggests there are a variety of genetic, environmental, health and lifestyle factors that play a role in the brain’s aging and cognitive capabilities as we age.^4^ Certain health and lifestyle factors that may potentiate various forms of cognitive impairments are modifiable risk factors (i.e., sedentation, hypertension, obesity, etc.) that can be mitigated by exercise.^3^ A 2019 meta-analysis observed significant associations linking exercise-induced improvements in physical and cognitive function.^5^

Regular exercise engagement throughout the entirety of one’s life may prove to be most protective against age-related cognitive decline (ARCD), with studies showing that higher rates of exercise from early to mid-adulthood likely reduce the risk of cognitive decline later in life.^6,7^ However, research also suggests that starting a regular exercise regimen later in life is still beneficial.^8^ Studies on older adult populations have found that participating in exercise programs, and increasing cardiorespiratory fitness, correlate with reductions in age-related neural changes ^9,10^ and improved cognitive performance.^5,8,11,12^ Reported benefits of cardiorespiratory (aerobic/endurance) exercise on the brain include improvements in executive function and divided attention,^13^ as well as specific executive function skills such as inhibitory control and working memory.^4^

In this study, we looked at the effects of high-intensity interval training (HIIT) on physical and cognitive performance in middle-aged male mice (18 months of age at study completion). Previously, we demonstrated HIIT can preserve physical function in adult (10m) and older adult (26m) mice.^14^ HIIT is a type of exercise performed in pre-determined intervals of higher-intensity that are interspersed with low-intensity (active rest) intervals. Thus, for the current study, we hypothesized there would be less cognitive and functional decline – as measured by the comprehensive functional assessment battery (CFAB) and a cognitive assessment battery (CAB) testing protocols – in the HIIT-exercised group (HIIT) as compared to the sedentary control group (SED). We measured exercise capacity and physical function with the previously-validated CFAB,^15^ and assessed cognitive function with a battery of commonly used cognitive/behavioral tests for mice (open field, y-maze, novel object recognition, and puzzle box).^16–21^ In addition, we determined the impact of HIIT on other potential markers of sarcopenia: body composition (EchoMRI), muscle wet weight, and maximal isometric plantar flexor torque. We observed significant changes between the two groups’ aerobic capacity and treadmill speed, but did not observe significant improvements in cognitive performance between or within groups.

## METHODS

### Subjects

We obtained C57BL/6J male mice (n=16) from The Jackson Laboratory at 10m (months) of age, equivalent to the start of early middle-age in humans.^14,22^ Mice began the training portion of the study at 14m with tissue collection for future use at 17m. Within a week of reception, before pre-testing began, one subject died of natural causes. Mice were humanely handled and treated under the approval of ECU Institutional Animal Care and Use Committee (IACUC). Mice were group-housed in 12-hour light/dark cycles at 22°C, with ad libitum access to food and water.

### Study Design

See **Figure 1** for study design. After an acclimation period, pre-intervention performance assessments (physical and cognitive) were completed. After pre-testing, mice were randomized into one of two groups: sedentary control (SED; n=7) and High-intensity Interval Training (HIIT; n=8). Randomization was followed by a 12-week HIIT training protocol with intervals based on a treadmill max speed test. The mice completed 6 weeks of training before they were re-tested on treadmill, after which the last 6 weeks of training commenced with an adjusted baseline for their intervals based on the retest.

**Figure 1.**
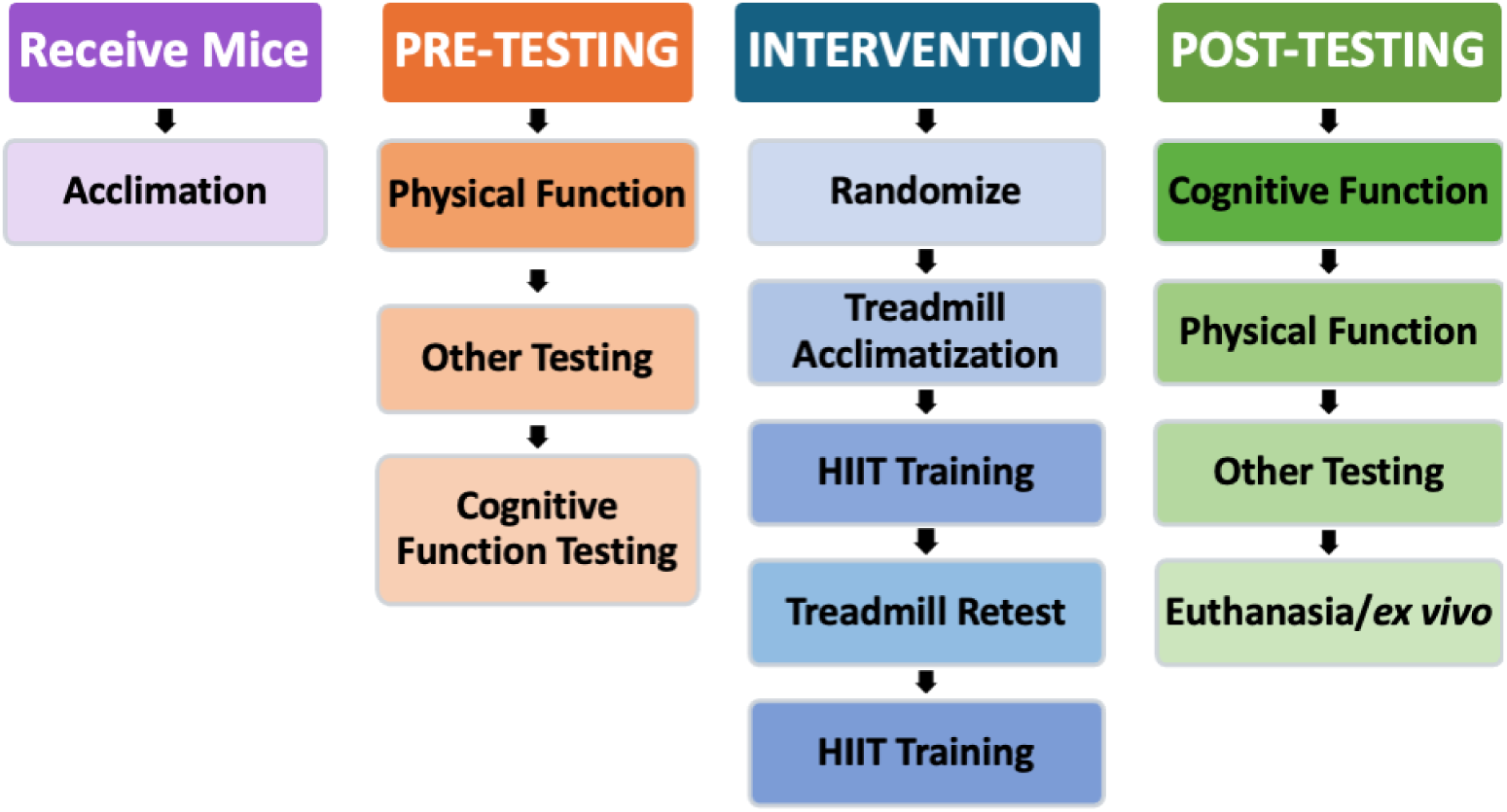
Study Design. After the initial acclimation period, the mice went straight into pre-testing. After baseline testing, the mice began the intervention stage of the experiment, where the HIIT group was acclimated to treadmill training for 1 week, trained for 6 weeks, retested for maximum speed on the treadmill, and then trained for another 6 weeks. Following the HIIT intervention, all mice were retested for physical and cognitive function. Physical and cognitive function testing were reversed between pre- and post-testing to account for any biases potentially created by testing in the same order.

### Intervention

After baseline testing and data analysis, mice were randomly assigned to one of two groups: high-intensity interval training (HIIT, n=8) or sedentary control (SED, n=7). Mice were group-housed in cages with environmental enhancements, such as a block or tunnel to play with, but no means of exercise (i.e., running wheels) available aside from general PA. Some mice needed to be individually housed due to over-aggression and fighting. The intervention consisted of one acclimation week, followed by 12 weeks of HIIT. Mice trained three times/week, one session every other day (i.e., Monday, Wednesday, Friday) to allow for rest and recovery between exercise sessions.

#### High Intensity Interval Training

##### Acclimation and Training

Before the first week of HIIT, mice were acclimated to the treadmill and interval speed changes for one week (3 sessions). One interval was run on the first day, two intervals on the second day, and three intervals on the third. By the first week of training, subjects were running three HIIT intervals at 75% of their cohort’s average maximum speed, transitioning to higher percentages and more intervals as the intervention progressed. Additional intervals were added as tolerated, up to a maximum of five, as the intensity/speed was also increased to the tolerance of the mice. If a mouse was unable to complete scheduled training at the expected intensity, it was relegated to the next slowest group until improvements were observed. After the sixth week (mid-point) of the intervention, mice were retested for a new max speed and their relative interval speeds were adjusted accordingly.

##### Exercise Sessions

The average max speed (Speedmax) of the mice, as measured by the baseline treadmill test, was used to determine percent max speed (%Speed_max_) for the HIIT intervals. Based on their fastest recorded speeds, mice were grouped into exercise cohorts, where cohorts of ≥ 2 animals with a similar Speed_max_ were exercised together. Subjects exercised 3 days/week with one rest day between training sessions (i.e., Monday, Wednesday, and Friday) and two rest days over weekends. Each HIIT session began with a 2-min warm-up at base speed (4 m/min), followed by 1-min intervals at sprint speed (with 30-second ramp-ups and 30-second ramp-downs, for a total of 2 minutes per interval), interspersed with a 1-min relative rest (walking speed). After the final HIIT interval, each session concluded with a 2-min cool-down at base speed.

##### Sham Treatment

As a sham treatment, SED mice were placed on the treadmill every day when we conducted HIIT training. The SED group did not exercise, but they were on the unmoving treadmill for the same amount of time as the HIIT session that day. Additionally, because HIIT mice were exposed to an active shock grid, the grid was also active when SED mice were on the treadmill for sham treatment. Since the HIIT group had opportunities to learn/gain experience from the training, the SED group needed to be allowed those same enrichment experiences to eliminate potential confounders.

### Performance Assessments

The same investigators performed the physical function and cognitive assessments throughout the study, with baseline and end-point assessments in the same order and at the same time of day. We used maintenance exercise training to maintain adaptations throughout the post-testing period.

### Physical Function Assessments

#### Functional Performance

Mouse physical function and exercise capacity was assessed via the Comprehensive Functional Assessment Battery (CFAB) pre- and post-intervention. We previously validated CFAB for male mice at three different ages (adult 6m, and older adult at 24 and 28m), and longitudinally in male and female mice over the lifespan ^15,23^. In summary, the mice were tested using a series of commonly-used well-validated determinants, including Grip Meter (forelimb strength), Inverted Cling (full-body strength/endurance), Voluntary Wheel Running (VWR; volitional exercise and individual activity levels), Rotarod (overall motor function, gait speed, balance/coordination, power generation), and Treadmill (maximum running speed and endurance/aerobic exercise capacity). CFAB was performed. Methodology for the determinants comprising CFAB have been previously described (21-25).

### CFAB Data Analysis

Traditionally, CFAB data analysis used a reference group of 6-month-old mice (mean and standard deviation; SD), where test results for each mouse are standardized (difference in SD from previously published six-month means) and summed together to quantify a composite CFAB score; a single numeric value representative of overall physical function capacity ^14,15,23,27^. In the current study, the baseline for standardization was the pre-intervention mean and SD of the entire sample (*n*=15), assessed prior to randomization, which was then compared to the individual mouse scores. The difference in pre- to post-testing was compared similar to the FIAV (frailty intervention assessment value) previously explained ^24^.

### Other Physical Measures

#### Body and Muscle Mass

We determined body composition (fat percentage, fat%) from pre- to post-intervention using an EchoMRI-700d. The EchoMRI-700^TM^ (Echo Medical Systems) is a quantitative nuclear magnetic resonance system that allows precise, whole-body composition measurements to be taken in live.

#### in vivo contractile physiology

We determined plantar flexor torque with an Aurora Whole Mouse 3-in-One physiology suite (Aurora Scientific), as described previously ^15,28^. Briefly, the mouse was anesthetized on a 37°C heated platform to maintain body temperature using ∼3% isoflurane with 1.5 liters/minute of O_2,_ to effect, from a VetEquip vaporizer nosecone to eliminate conscious control of skeletal muscles. We positioned the knee at 90° with the tibia parallel to the platform and then clamped the femur lateral and medial epicondyles in place to prevent the leg from shifting yet allowing for free movement below the knee. The foot was set into a footplate connected to a force transducer, and the height adjusted to firmly set the heel into the bottom of the plate. Using needle electrodes placed subcutaneously, we determined the optimal location and current needed to produce a maximum torque twitch. This current and needle placement was maintained during a torque/frequency curve (a single pulse, then 10, 40, 80, 100, 120, 150, 180, and 200 Hz) to find maximum tetanic isometric torque of the plantar flexors.

### Cognitive Function Assessments

#### Cognitive Performance

We assessed cognitive function with our Cognitive Assessment Battery (CAB) pre- and post-intervention. Cognitive performance was determined through the application of memory, behavioral and executive function tasks. The tests included Open Field (anxiety, locomotor and exploratory behavior)^20,29^, Y-maze (exploratory behavior and spatial working memory)^18^, Novel Object Recognition (exploratory behavior and long-term memory)^20,29^, and Puzzle Box (short-term memory and executive function)^16,19^. All cognitive/behavioral assessments were video recorded with a GoPro HERO6 Black for later analysis and data quantification. The CAB outcome measures were analyzed individually in this study. Individual testing procedures are discussed in more detail in the Supplemental Methods, but in brief:

#### Open Field

Open Field (OF) is a commonly used behavioral test for assessing general locomotor activity, anxiety levels, and exploratory behavior in mice ^20,29^. The testing arena was a 58x58x40cm box, made of a non-abrasive plastic, with an open top for direct lighting and video recording. Before testing, we assessed light distribution using Light Meter LM-3000 to ensure uniformity of brightness (750 ± 10 lux) across the testing arena. We applied direct lighting with a single LED lamp positioned over the center of the arena. The outcome measures for OF were the number of entries into the center and time (s) spent in the perimeter.

#### Y-maze

Y-maze assessed spatial working memory, as described in the literature ^18^. We used a custom-built symmetrical Y-shaped maze with a non-reflective, neutral-colored surface (beige), details in Supplemental Methods. In brief, each mouse was positioned into any one arm (designated Arm A) facing toward the maze’s center and were then allotted 8 minutes of uninterrupted exploration. Entry into an arm was defined by the animal having all four paws inside the arm. Total entries were counted as an indicator of locomotor activity, whilst any latency of a mouse leaving its starting arm was an indication of emotionality-related behavior. The outcome measure was the percentage of spontaneous alternation performance (%SAP). The criteria required to be considered a spontaneous alternation (SA) in this experiment was defined as sequential entries into all three arms in overlapping triplet sets (i.e., ABC, BCA, CAB, or vice vera) as shown in **Figure S3**. The %SAP was defined as the ratio of total alternations to possible alternations (%SAP = SA / [total arm entries – 2] x 100).

#### Novel Object Recognition

We administered the Novel Object Recognition (NOR) test to assess long-term memory and exploratory behavior in mice ^20,29^. Discrimination of novel versus familiar stimuli requires intact perceptual systems. Therefore, if a mouse spends more time exploring a novel object (NO) compared to a familiar object (FO), it is indicative of an intact memory ^29^. A discrimination ratio (DR) was calculated to quantify novelty preference by subtracting the time (s) spent exploring the FO from time (s) spent exploring the NO and dividing that by the total object exploration time (s; DR = (NO – FO) / total exploration). The outcome measure was the discrimination ratio.

#### Puzzle Box

The puzzle box test was designed to assess executive function skills in mice via working memory and problem-solving requirements. The test was adapted from previously used versions ^16,19^. For this adaptation of the puzzle box assessment, a PVC pipe connects the big OF arena to a much smaller “puzzle box” (17.3×21×17.6cm). The OF arena was the open, brightly illuminated starting area, while the smaller, darker puzzle box was the objective area. To access the objective area, subjects must climb into the tunnel and make their way across.

Puzzle box tasks began with each mouse positioned in the center of the wall directly across from the puzzle box access, and assessed on how long it took to complete the objective. A treat/prize (i.e., unsalted walnut, almond, or plain Cheerio) was used as an additional incentive. With each test, we modified the access point to increase difficulty of reaching the puzzle box; the mice encountered various challenges designed to increase difficulty of accessing the tunnel/puzzle box. We tested the mice without any obstruction to accessing the puzzle box (Day 1), then we added an obstruction blocking the exit of the PVC pipe that the mice had to simply knock down to gain access to the puzzle box (Day 2). The last task had two trials (T1 and T2), and for each trial the entrance point was facing a different direction (Day 3).

### Statistical Analysis

Independent samples t-tests were used to compare mean differences between the HIIT and SED groups performance scores on CFAB and CAB assessments. Paired samples t-tests compared changes within the groups. CFAB functional determinants were assessed using 2×2 mixed-design ANOVA (see results for more details). Differences were deemed significant at p<0.05. Data expressed as mean ± SE (standard error), unless otherwise indicated.

## RESULTS

We used student’s independent samples, paired t-tests, and 2×2 ANOVA, as appropriate, to compare dependent variables with the results reported in the appropriate tables, alongside mean, SD, SEM, effect size (Cohen’s D or η^2^) skew, kurtosis, and the results of the Kolmogorov-Smirnov and Shapiro-Wilks tests for normality. See **Online Only Supplemental Datasets S1-S6** for further details.

### Physical Function (CFAB): Improved Aerobic Capacity from HIIT

We determined physical function and exercise capacity, pre- and post-intervention using CFAB. Note that all CFAB determinants were normally distributed, as we have previously determined and published ^15^, with the exception of inverted cling which we transformed to log10 to meet the criteria of normality for our statistical tests.

There were no significant changes (2×2 mixed ANOVA [2 groups: SED and HIIT; and 2 time points: pre- and post-intervention]) in grip meter (strength), inverted cling (overall strength/endurance), voluntary wheel running (volitional exercise), or rotarod (overall motor function) with training either between or within subjects compared to sedentary mice. See **Dataset S1 and Figure 2** for more details.

**Figure 2.**
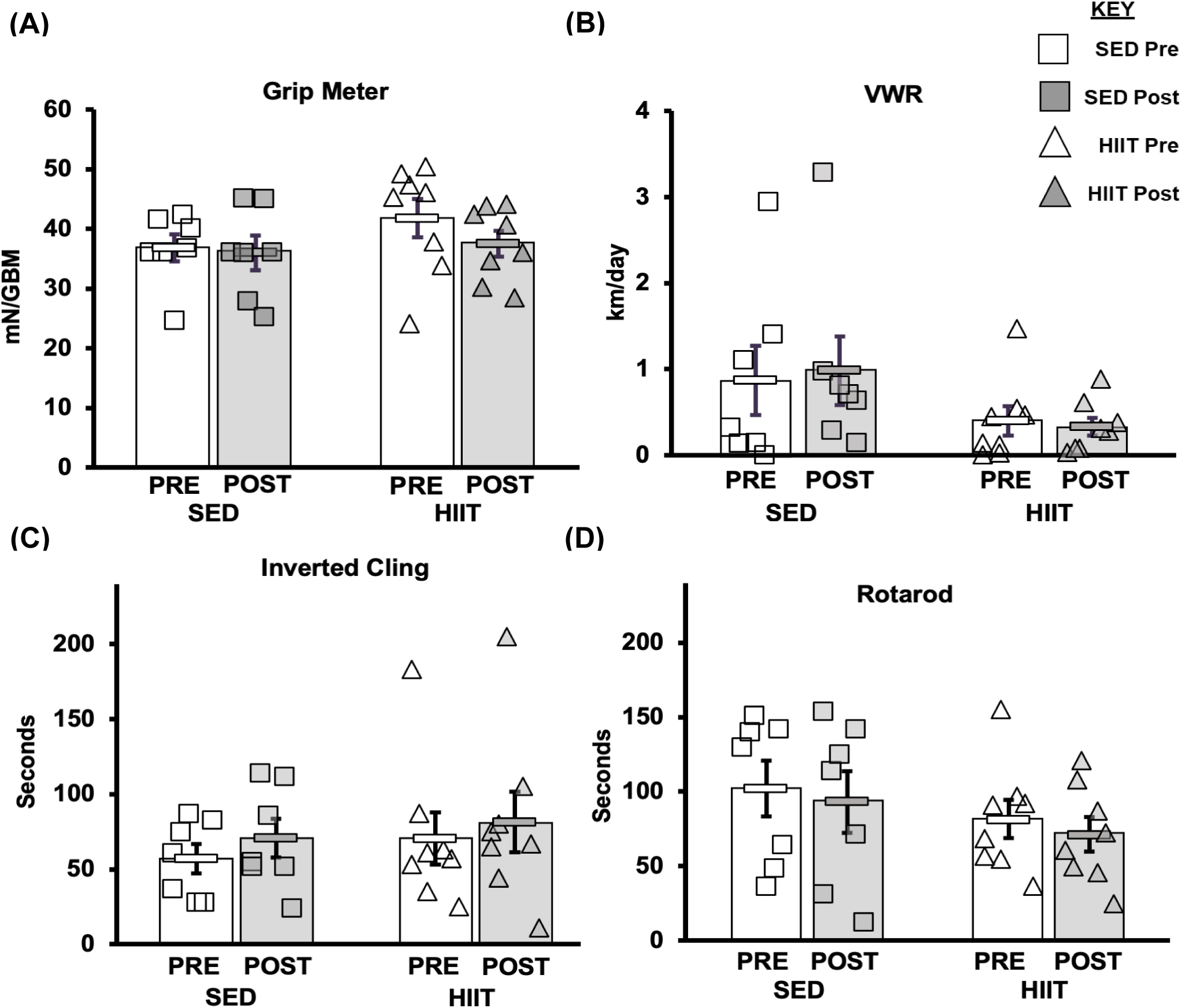
CFAB Determinants. **A) Grip Meter. B) Voluntary Wheel Running (VWR), C) Inverted Cling, D) Rotarod**. From pre- to post-intervention none of these tests demonstrated significant training effects. **KEY:** mN/GBM = milliNewton per gram of body mass. SED = sedentary control group, HIIT = high intensity interval training group, PRE = baseline before intervention period, POST = value after the intervention period.

To assess changes in aerobic capacity and running speed, the treadmill test was administered pre- and post-intervention for both groups. For the HIIT group, treadmill time significantly increased by 28.0% pre- to post-training, while the SED group declined in performance by 41.4% (within-subjects effects of time*groups F=21.381, p<0.001, partial η^2^=0.622; between-subjects effects of time*groups F=5.572, p=0.035, partial η^2^=0.300).The HIIT group increased a mean of 138.1 seconds and the SED group declined a mean of 217.7 seconds (see **Dataset S1 and Figure 3**) from pre- to post-intervention (post-hoc testing by independent samples t-test t=5.572, p<0.001; within-subjects changes were both significant and had large effect sizes: paired samples t-test SED t=3.901, p<0.008, Cohen’s d=-1.475; HIIT t=2.612, p=0.035, Cohen’s d=0.923). In addition, the average maximum speed from pre- to post-training (see **Figure 4**) significantly changed. Overall physical function as measured with CFAB did not alter pre- to post-training (within-subjects effects of time*groups F=0.070, p=0.795, partial η^2^=0.005; between-subjects effects of time*groups F=0.037, p=0.851, partial η^2^=0.003). See **Figure S1** and **Dataset S1** for further details. At pre-intervention testing, there was no difference observe between groups (p=0.555). However, the HIIT group significantly increased (p=0.038) in average maximum speed pre- to post-intervention while performing significantly better (p = 0.00008) than SED which decreased (p=0.019).

**Figure 3.**
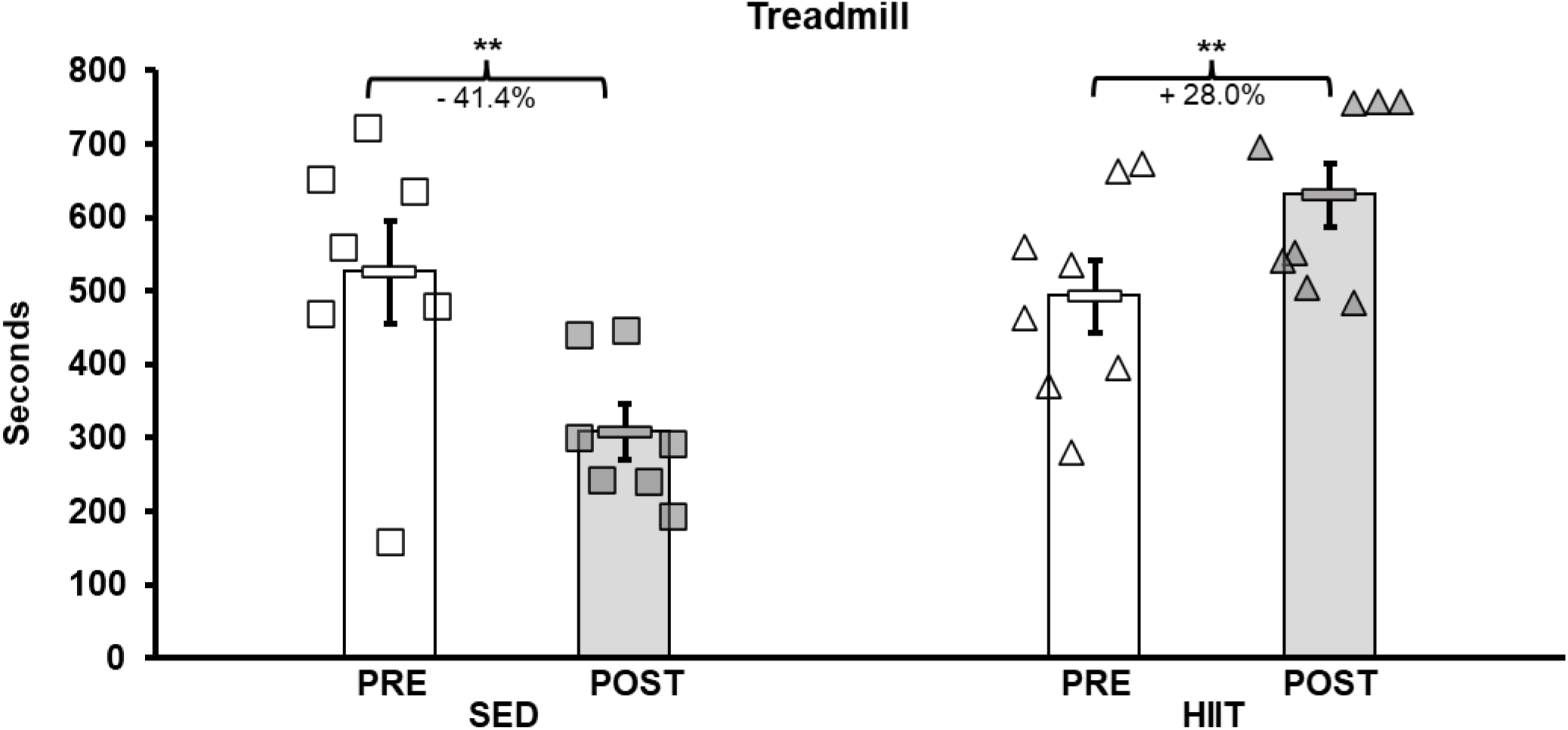
Treadmill Max Speed Test. The mice were assessed pre- and post-intervention for their aerobic capacity. Pre-intervention testing revealed no significant difference between the two groups’ aerobic capacity, while post testing showed significant differences between groups and within each group’s pre- to post-intervention aerobic capacity. Significant decrease in treadmill time for the SED group (-217.7 seconds), and a significant increase in treadmill time for the HIIT group (+138.1 seconds). KEY: SED = sedentary control group, HIIT = high intensity interval training group, PRE = baseline before intervention period, POST = value after the intervention period, **=P<0.01.

**Figure 4.**
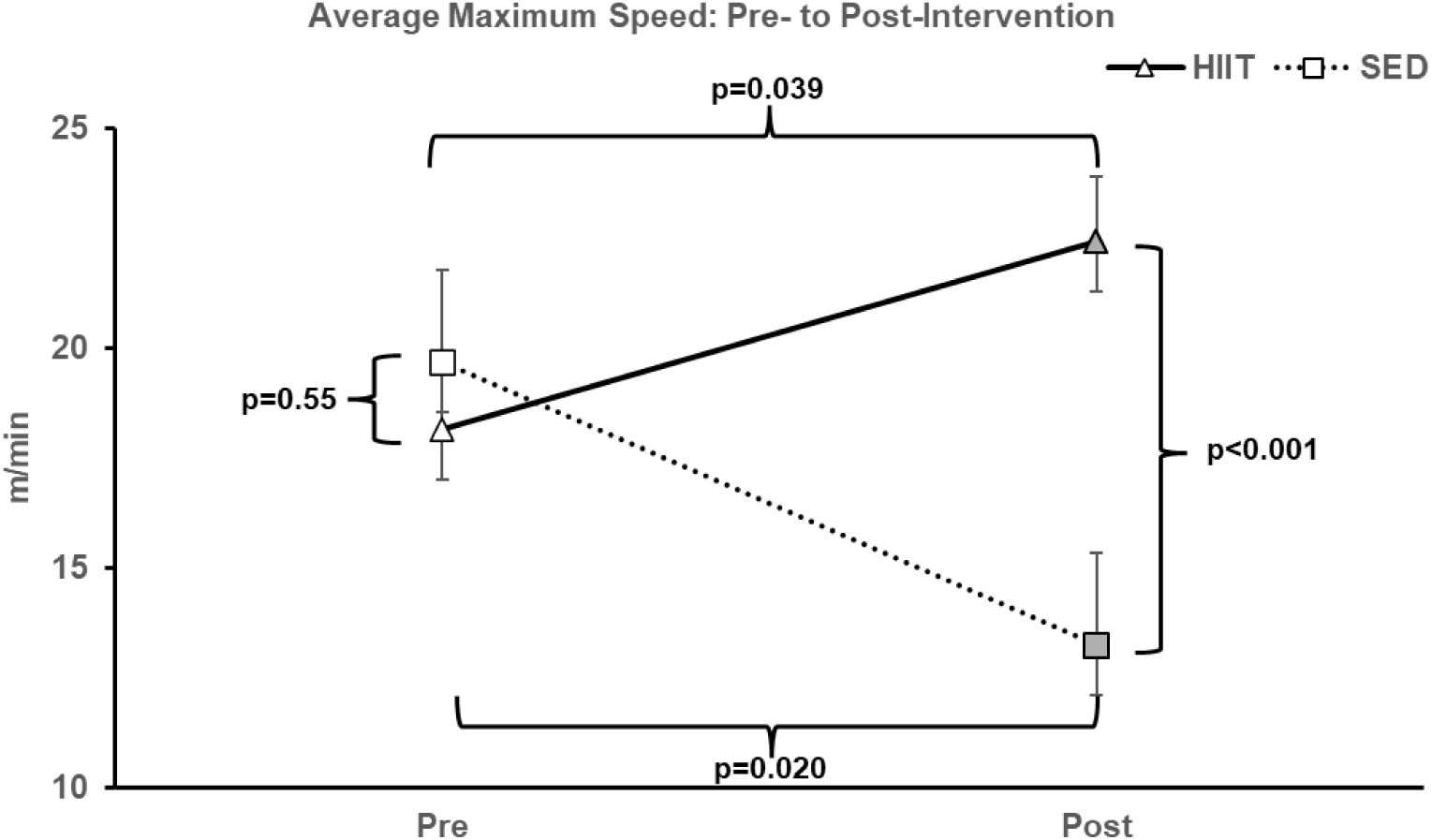
Average maximum treadmill speed. At pre-intervention testing, there was no significance observed between groups. However, after the training period the HIIT significantly increased within group and compared to SED (decreased significantly within group). KEY: SED = sedentary control group, HIIT = high intensity interval training group, PRE = baseline before intervention period, POST = value after the intervention period.

### Other Measurements

#### Body Composition

Body mass (grams; g) was measured weekly throughout intervention (see **Figure 5**), prior to each EchoMRI, and at euthanasia. There was no difference in any of the measurements prior to training. The difference between body mass, fat, and fat% measured at the time of EchoMRI was significantly larger within groups from pre- to post-training (2×2 Repeated Measures ANOVA: F=20.062, 46.845, and 35.899, respectively; all p<0.001). Between subjects’ lean mass tended to decrease in SED and remain the same in HIIT from pre- to post-training (2×2 Repeated Measures ANOVA: F=3.551, p=0.082); although, while the lean mass difference between groups had a strong effect size (-0.746), it was not significantly different (independent samples t-test, and p=0.173). See **Dataset S2** and **Figure 5** for more details.

**Figure 5.**
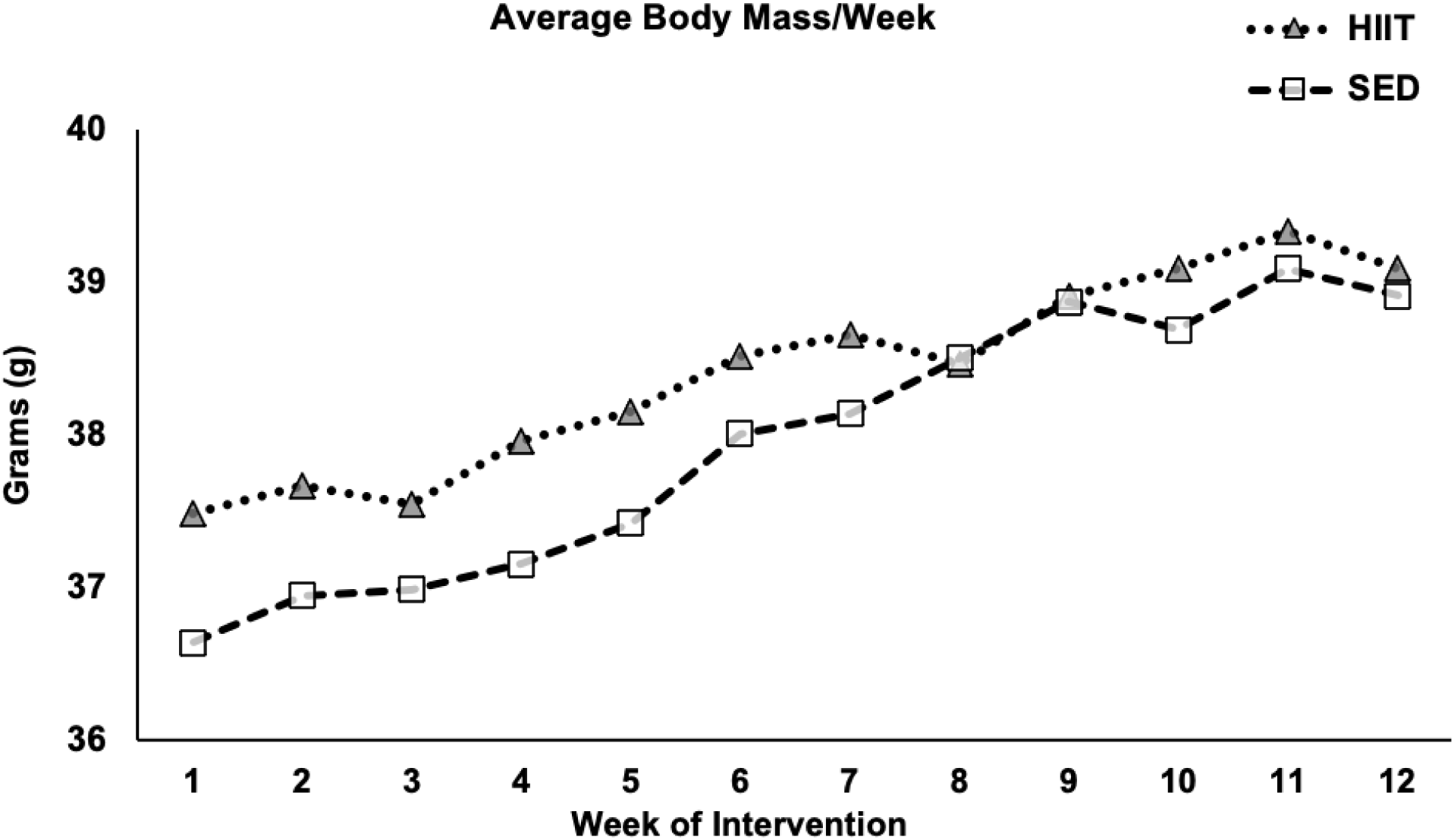
Average body mass. The mice were exercised or exposed to sham treatment 3 days/week. Every week, before their third session, the mice were weighed. The figure above shows the average body mass for each group (HIIT and SED) for each week of the intervention. KEY: SED = sedentary control group, HIIT = high intensity interval training group.

### Muscle Mass

The gastrocnemius (GAS), plantaris (Plant), tibialis anterior (TA), extensor digitorum longus (EDL), soleus (SOL), and heart muscles were collected, blotted dry, and weighed (grams; g) following euthanasia. Between-groups analysis (independent samples t-test) showed no significant difference for each muscle mass or total muscle mass. See **Dataset S3** for more details.

#### *in vivo* Contractile Physiology

Maximum isometric plantar flexor torque was measured pre- and post-training. There was no difference in groups prior to training. We found no significant change (paired samples t-test) in either maximum torque (mN*m) or normalized torque (mN*m/grams of body mass; paired samples t-test SED t=1.369, p=0.243, Cohen’s d=0.612; HIIT t=0.17, p=0.873, Cohen’s d=0.076) following training. See **Table S4** for more details.

### Cognitive Function (CAB)

We measured different parameters of cognitive function using four different tests, pre- and post-intervention, and performed means testing with independent samples t-tests between groups and paired samples t-tests within groups. See **Table S5** and supplemental figures for further details.

#### Open Field

For the OF test, entries into the center and time spent in the perimeter were measured, with the HIIT group spending significantly more time in the perimeter (p=0.016), which is indicative of greater anxiety-related behavior. The SED group showed trends toward a similar result, with fewer center entries and more time spent in the perimeter zone, but nothing of significance. However, there were limited defecation and urination events during, both pre-(n=4 total for all mice) and post-testing (none), indicating little evidence of anxiety in this regard. See **Figure S2**.

#### Y-Maze

The Y-maze assesses subjects’ spatial reference memory by measuring the number of Spontaneous Alternations (SAs) made in the allotted test time. SAs were converted into the percentage of the total number of alternations. The number of spontaneous alternations increased significantly in both groups (HIIT, p=0.002; SED, p=0.013) from pre to post. However, total arm entries also increased significantly in both groups (HIIT, p=0.001; SED, p=0.007), thus resulting in no significant improvements in %SAP. Although total arm entries were greater for both groups, the increase was still more evident in the HIIT group (see **Figure S3**).

#### Novel Object Recognition

There were no significant changes in NOR (p=0.757) from pre- to post-training. See **Figure S4**.

#### Puzzle Box

There were no significant changes in the puzzle box tests (Blocked Exit, p=0.689; Different Entrances, p=0.378). However, for the blocked exit task, between-groups analysis showed the SED group was significantly faster at entering the tunnel (p=0.036) pre-intervention but could not replicate that significance in post-testing (p=0.306; HIIT increased a mean of 17 seconds; SED declined a mean of 39.8 seconds from pre- to post-intervention). There was also no significance observed in the blocked exit task pre-intervention, but the SED group was significantly faster than HIIT at removing the obstruction (p<0.001) to the puzzle box during post-testing. Though between-groups analysis did show significance for other test variables collected, none fit normality. For example, in the differing entrances task, analyses showed the SED group did significantly better finding their way into the second (T2) entrance (p=0.005) pre-intervention but did not prove to be significantly better in the post assessment (p=0.904). See **Figure S5**.

### Exercise Intensity/Work

#### HIIT Intensity and Work Increased Through Middle Age

Throughout the intervention, the HIIT group’s exercise intensity and work performed increased relative to how each mouse was responding to their current load. Over the course of the HIIT intervention, exercise intensity (%Speed_max_) increased an average of 14.36%. Between the midway point (week 6) treadmill retest and the last day of training (week 12), exercise intensity increased an average of 15.27%.

We calculated the total amount of work (m*gbm, meters*grams of body mass) for each HIIT session performed. The average significant difference (p<0.05) in power produced (work performed per minute) between each subject’s first and last HIIT session was equal to 40.32 (m*g)/min. This indicates exercise adaptation.

## DISCUSSION

The current study was designed to determine the effects of High-Intensity Interval Training (HIIT) on functional and cognitive performance in a group of middle-aged C57BL/6 male mice compared to a sedentary control group (SED), pre to post a 12-week treadmill HIIT protocol.^14^ Overall CFAB did not change with exercise in this population, though we observed significant improvements in the HIIT group’s aerobic capacity and treadmill speeds with a corresponding decline in the sedentary mice. For body composition, we observed significant increases pre- to post-training between body mass, fat, and fat % within groups, and a strong effect size for lean mass difference between groups but it was not significant. Finally, we did not detect any significant cognitive changes between the two groups.

### HITT

In recent years, HIIT has been popularized as a safe and time-conscious alternative mode of exercise. HIIT exercise has been reported to have positive effects among several chronic diseases ^30–33^ often associated with cognitive decline/diseases in older adults.^34^ Furthermore, even short-term (6 weeks) HIIT has been shown to produce physiological and physical fitness improvements similar to, and sometimes better than, endurance and/or resistance training in middle-aged men.^35^

Our 12-week treadmill HIIT protocol was adapted from previous studies.^14,36^ While Seldeen et al. (2018) used mice 22m of age, the findings from Pajski et al. (2024) were based on mice aged 6m–10m and 22m–26m. Thus, there remained a gap in the literature for the 10m–17m (middle age) age gap.

Graber et al. (2021) validated the use of CFAB in male mice at 6m, 24m, and 28+m of age, observing overall age-related physical function decline. In the current study, CFAB, administered pre- and post-intervention (10m and 17m old, respectively), detected no significant changes in the intervention assessment value (IAV) determinants of grip meter, inverted cling, VWR, rotarod, or overall. This contrasts with prior findings in which significant improvements in, or preservation of, physical function were observed.^14,36,37^ However, Seldeen and colleagues did not observe improvements in rotarod for the HIIT group in their 2018 study, nor did they observe significant changes in rotarod or inverted cling for either group (SED or HIIT) in their 2019 study.^36,37^ Notably, these studies investigated different age groups than the current study. In recent work, a relative plateau in functional capacity between 12m - 18m in male C57BL/6 found by Pajski et al. (2024) indicated a stability of function, instead of decline, in early- to later-middle-age which might partly explain our results. Although CFAB did not improve overall with training, there was a marked improvement in aerobic capacity (treadmill time) and treadmill speeds, while the sedentary control mice declined. This finding is consistent prior studies ^14,35–37^ in both animals and humans.

Contrary to our hypothesis, neither group declined in cognitive function and exercise did not result in improvement. In the literature, there appears to be a correlation between age and cognitive performance in C57BL/6 male mice, with statistical significance observed in the younger and older age groups.^38,39^ However, it is unclear whether the studies observed significance between the 12m—20m age group. Thus, age-related cognitive function patterns during normal aging in BL/6 mice need further study.

In human research, there are mixed reviews on whether exercise, or specific types of exercise, have any significant effect on cognition. A Gates et al. (2013) review of exercise effect on cognitive function in older adults (65-95 years old) with mild cognitive impairment (MCI), revealed little evidence to support exercise-induced functional improvement.^40^ However, in 2022, results from a comparison study on the effects of HIIT and moderate-intensity continuous training (MICT) on cognitive function indicated exercise alone could promote cognitive function independent of the exercise type.^41^ A meta-analysis by Cammisuli et al. (2017) reported aerobic exercise as beneficial for patients with mild cognitive impairment (MCI), but they observed only moderate effects of aerobic training on global cognition, inhibitory control, logical memory, and divided attention.^42^ Subsequently, Cammisuli and colleagues (2018) reported little evidence for improvements in AD patients’ cognition from aerobic exercise.^43^ However, other reviews and meta-analyses present evidence of the efficacy of exercise training to improve cognition, with resistance training often observed as a superior intervention in some cognitive domains and relative exercise intensity playing a central role—though very little information about HIIT is available.^44–47^

The literature suggests exercise intensity may be a critical factor in maintaining/improving memory, with higher intensities reportedly improving memory in sedentary older adults.^48^ Contrary to our findings, where significant aerobic capacity improvements in the HIIT group showed no cognitive improvements, Kovacevic et al. (2020) found a significant correlation between increased cardiorespiratory fitness and memory improvements in humans >60 years of age.^48^ However, our mice were only the equivalent to that of early 50’s in human years by study’s end.

### Exercise and Body Composition

For body composition, we observed significant increases between body mass, fat, and fat % within groups from pre- to post-training, and a strong effect size for lean mass difference between groups but it was not significant. Using older-adult (24m) mice, Seldeen et al. (2018) reported significant declines in fat % for their SED mice, while their HIIT group exhibited no such decline, maintaining greater fat % than the control group. Previously, we reported marked fat% declines in both 26m exercise groups (voluntary wheel running and HIIT) compared to increases in controls while 10m groups all increased, though increased fat gain was mitigated significantly with exercise.^14^ Natural body composition patterns seen in humans and rodents may explain the findings of Seldeen et al. (2018), Pajski et al. (2024) and the current study. A recent review found that, on average, peak fat mass in mice – provided food ad libitum – occurs between 12m and 24m (roughly 40-80 years in humans), while fat mass decline is observed between 17m and 24m (57-80 years in humans).^49^ As such, it was reasonable for our mice to continue gaining fat regardless of training or sedentation.

### CAVEATS

One limitation of the current study is that due to within-house fighting, certain mice in both groups required individual housing due to fighting, while the rest remained group-housed. According to a meta-analysis reviewing common methodological issues in animal research investigating the effects of exercise on cognition, singly-housed rodents may suffer from social isolation.^50^ The study found that social isolation was associated with a greater effect of exercise on cognitive performance. Due to randomization, 42.85% of the SED mice were singly housed, while just 25% of the HIIT mice were singly housed. This could potentially be related to the lack of cognitive differences observed between groups. Additionally, this study only had male mice of a single strain and age. Future studies will determine the effects of HIIT on cognition in older cohorts and determine any sexual dimorphisms.

## CONCLUSION

We did not observe statistically significant changes between groups, pre- to post-intervention, in grip meter, inverted cling, VWR, rotarod, or overall IAV (ΔCFAB). However, the HIIT group significantly increased in aerobic capacity and treadmill time, while the SED significantly declined. No significant between-group differences were observed for any of the cognitive measures assessed, pre to post. The lack of cognitive changes observed could be due to the age of the mice, as they may not have started experiencing any age-related cognitive decline yet, which would mitigate any potential effects of the HIIT intervention. More research is needed to determine the interactive effects of exercise and cognition across a broader range of aged mice and whether biological factors such as sex and strain contribute to these differences.

## Supporting information

Online Only Supplement

## Acknowledgements

**Animal Use Statement:** All animals were treated humanely under East Carolina University IACUC guidelines.

## Author Contributions: in order of contribution, * = equal contribution

The authors recognize the relative contributions as: Conceptualization, TGG, JS; Methodology, JS, TGG, TT; Validation, TGG, JS; Formal Analysis, JS, TGG; Investigation JS, TGG; Resources TGG; Writing – Original Draft JS,TGG; Writing – Review & Editing JS,TGG,TT; Supervision TGG; Project Administration TGG, JS; Funding Acquisition TGG.

## Acknowledgements

The authors wish to acknowledge Brandon Baucomb for technical assistance. We also acknowledge the East Carolina Obesity and Diabetes Institute for their role in providing resources and equipment for this study (e.g., EchoMRI).

