## Supplementary material for "Effects of High-Intensity Interval Training on Physical and Cognitive Function in Middle-Aged Male Mice": Online Only Supplement

### Online Only Supplemental Section for: Effects of High-Intensity Interval Training on Physical and Cognitive Function in Middle-Aged Male Mice

Justin C. Stephenson[1*], Tuan D. Tran[2], Ted G. Graber[1,2,3,4,5*]
Affiliations: East Carolina University (ECU) [1]Dept. of Kinesiology, [2]Dept. of Psychology, [3]Dept. of Physical Therapy, [4]Dept. of Physiology, [5]East Carolina Obesity and Diabetes Institute

[*]Corresponding/Senior Author:
Ted G. Graber, PhD.
ECU College of Allied Health Sciences
Department of Physical Therapy
Health Sciences Building, Rm 2410 | Mail Stop 668
Greenville, NC 27834


**Available on FigShare @ doi: 10.6084/m9.figshare.28410149**

**Table of Contents:**

| Online Item | Title | Location |
|---|---|---|
| **Figure S1** | **Total CFAB Scores and Intervention Assessment Values** | **Page S2** |
| **Figure S2** | **Open Field Pre- to Post-Training** | **Page S3** |
| **Figure S3** | **Y-maze Pre- to Post-Training** | **Page S4** |
| **Figure S4** | **Novel Object Recognition Pre- to Post-Training** | **Page S5** |
| **Figure S5** | **Puzzle Box Pre- to Post-Training** | **Page S6** |
| **Dataset S1** | **Physical Function** | **Excel File** |
| **Dataset S2** | **Body Composition** | **Excel File** |
| **Dataset S3** | **Muscle Mass** | **Excel File** |
| **Dataset S4** | **Contractile Function** | **Excel File** |
| **Dataset S5** | **Cognitive Function** | **Excel File** |
| **Dataset S6** | **Exercise Intensity/Work** | **Excel File** |

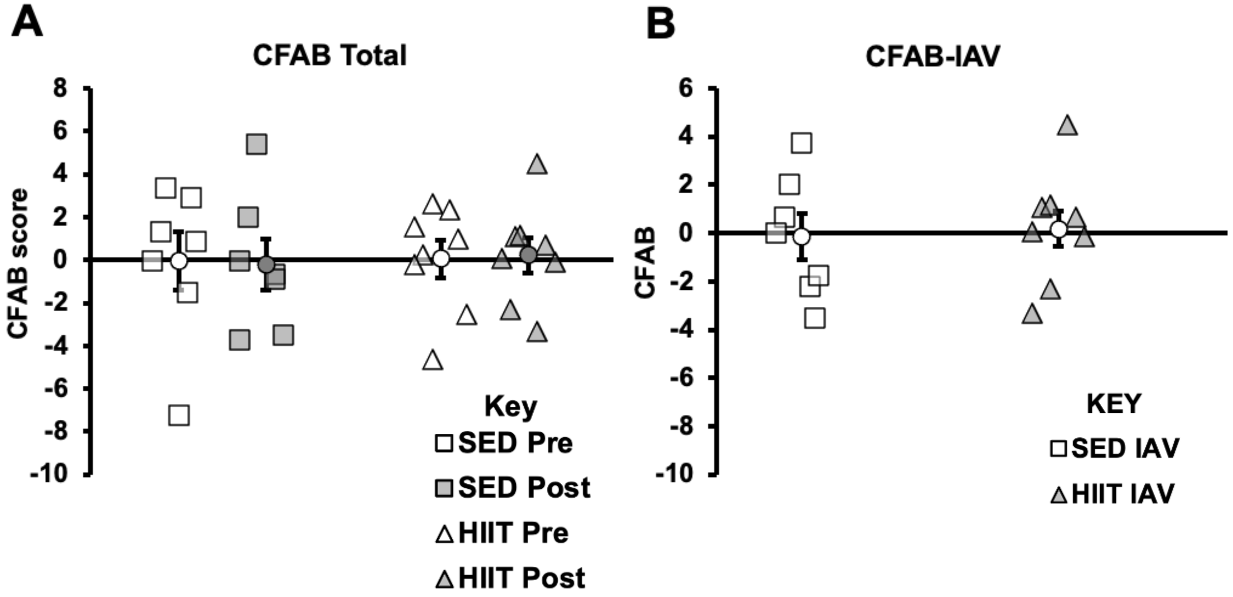


**Figure S1: Total CFAB Scores and Intervention Assessment Values.** CFAB analysis uses a reference group of 6-month-old mice (mean and standard deviation; SD), where test values are standardized via the distance of each individual mouse's score from the 6m mean. Distance from that mean is measured in units of the reference group's SD. The standardized scores for each functional test are then added together to form the CFAB composite score: a single numeric value representative of overall physical function capacity (A). An intervention assessment value (B) was also used for analysis in this study, in which standardization was achieved by using baseline mean and SD of the entire sample (n = 15) before randomization occurred.

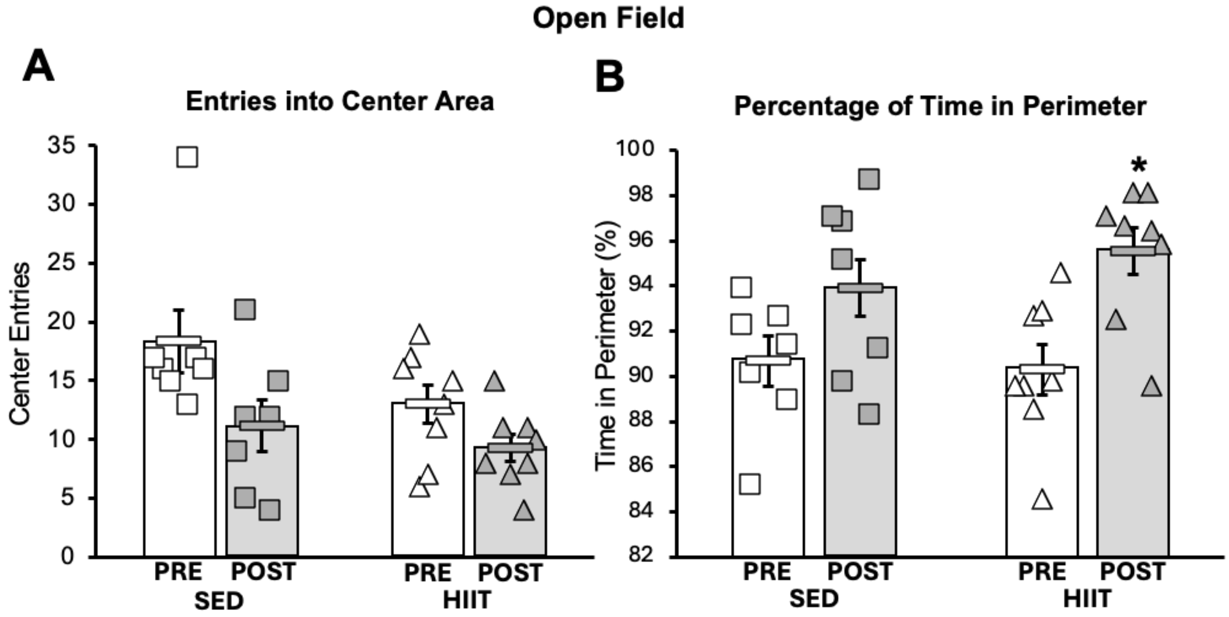


**Figure S2: Open Field Pre- to Post-Training.** The number of times a mouse entered the center area of the open field box in the allotted time is shown on the left (A). The percentage of their allotted time spent in the perimeter region of the box is shown on the right (B). Naturally, as center entries (A) decrease, the amount of time spent in the perimeter (B) increases. However, this trend of lessened time in the center and added time in the perimeter is indicative of increasing anxiety from pre to post.

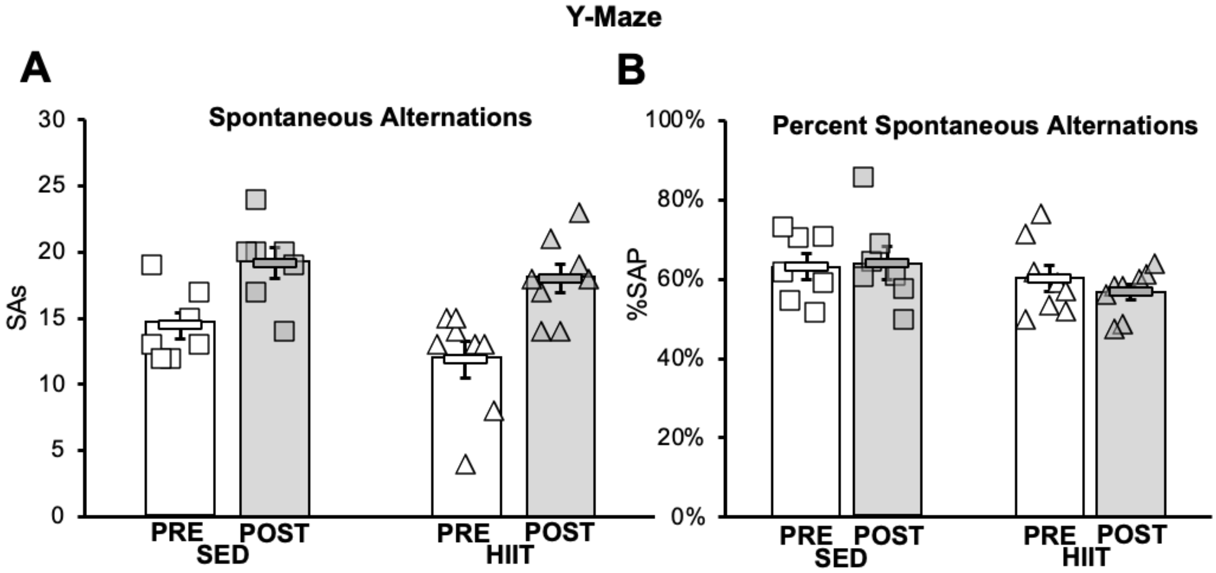


**Figure S3: Y-maze Pre- to Post-Training.** To assess spatial working memory with Y-maze, spontaneous alternations (SAs) were counted (left). Though the number of SAs increased more for the HIIT group (+49 overall) compared to SED (+33 overall), the percentage of SA performance (right; %SAP=SAs/[total entries-2]) showed no significant differences. %SAP decreased for HIIT (-3.40%) and remained relatively the same for SED (+0.94%).

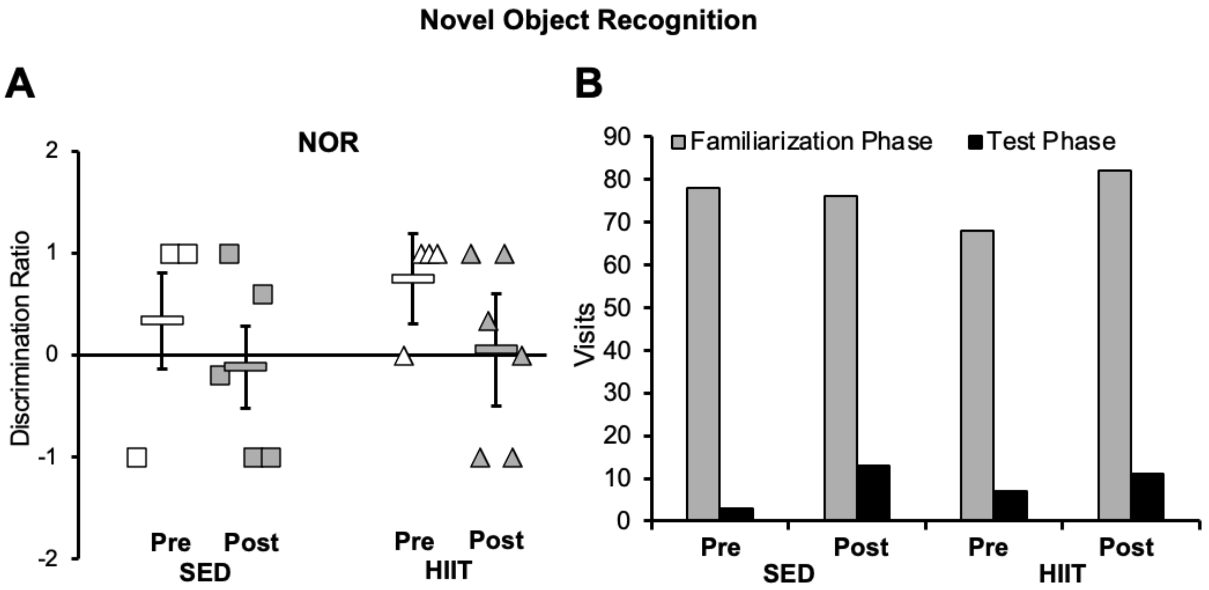


**Figure S4: Novel Object Recognition Pre- to Post-Training.** The discrimination ratio (DR) is a quantification of the between-object bias observed during the NOR test (A). It is calculated as DR=(Novel Object visits – Familiar Object visits) / total visits. The amount of exploration is the number of visits to either object (B). This was significantly reduced across the board between the familiarization phase and the testing phase, indicating the mice may have lost interest in the objects after the familiarization phase, thus, explaining the DR results (A).

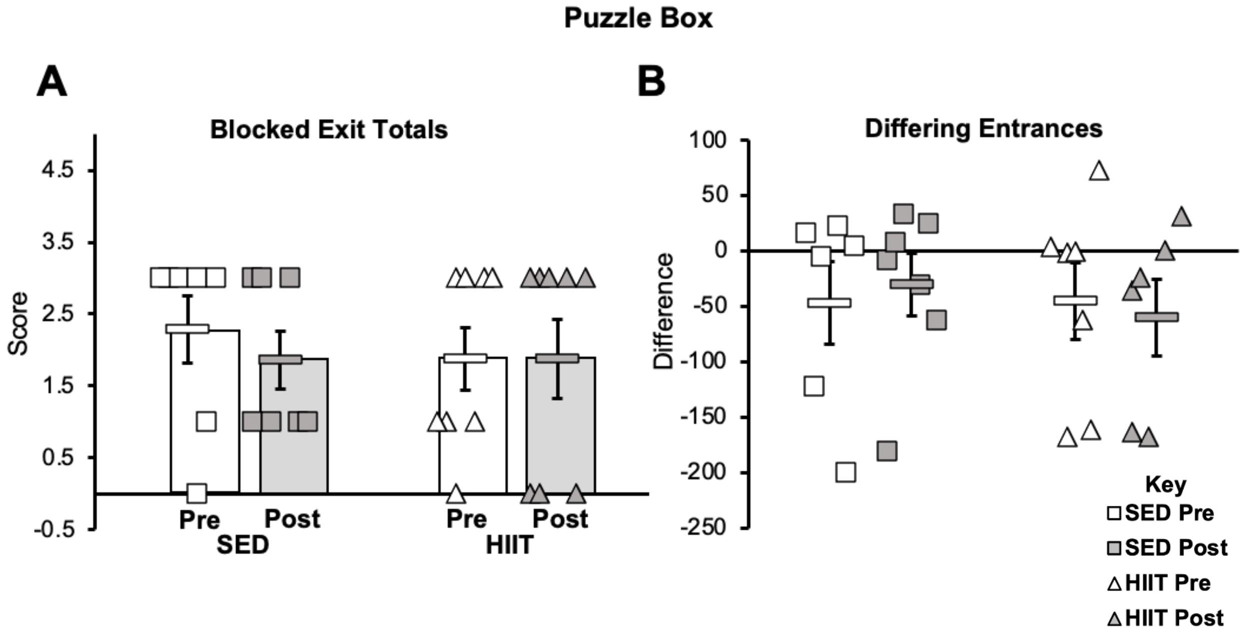


**Figure S5: Puzzle Box Pre- to Post-Training.** Though no significance was observed, the blocked exit test (A) showed no change in the HIIT group's ability to problem solve, while the SED group did decline in this area. The differing entrances test (B) was analyzed by finding the difference between trial 1 and trial 2, then calculating the difference of those differences from pre to post. Meaning, the more negative from pre to post, the faster the group solved the task.
